# High-resolution spatial profiling identifies disease-specific molecular architecture in palmoplantar pustulosis

**DOI:** 10.64898/2026.05.08.723901

**Authors:** Kazuki Yatsuzuka, Jun Muto, Yoichi Mizukami, Keishiro Isayama, Daisuke Shiokawa, Mei Miyazaki, Teruko Tsuda, Ken Shiraishi, Yasuhiro Fujisawa, Masamoto Murakami

## Abstract

Palmoplantar pustulosis (PPP) and dyshidrotic eczema (DE) are chronic vesiculopustular dermatoses with overlapping clinical presentations but distinct underlying biology. Although comparative transcriptomic and proteomic analyses between PPP and DE have been reported, they remain limited in number and scope, with no comprehensive understanding of their distinct molecular signatures. Moreover, their molecular mechanisms remain unclear, and currently available therapeutic options are limited. To clarify disease-specific epidermal programs underlying vesicle formation, we conducted Visium HD spatial transcriptomic analysis of FFPE lesional skin samples obtained from patients with PPP and DE, followed by immunohistochemical validation against normal palmoplantar skin controls. Spatial clustering identified a keratinocyte subpopulation adjacent to vesicles that exhibited distinct transcriptional programs in the two diseases. In PPP, vesicle-associated keratinocytes demonstrated marked downregulation of aquaporin-3 (AQP3) and E-cadherin, together with strong, spatially localized activation of JAK-STAT3 signaling. Conversely, DE exhibited diffuse AQP3 expression and more homogeneous activation of JAK-STAT3 signaling throughout the epidermis. These results indicate that, although PPP and DE share inflammatory pathways, they differ substantially in their spatial molecular architecture. Reduced AQP3 expression and localized STAT3 activation may contribute to vesicle formation in PPP, supporting our previous hypothesis that implicates intraepidermal sweat leakage as a pathogenic mechanism in PPP.

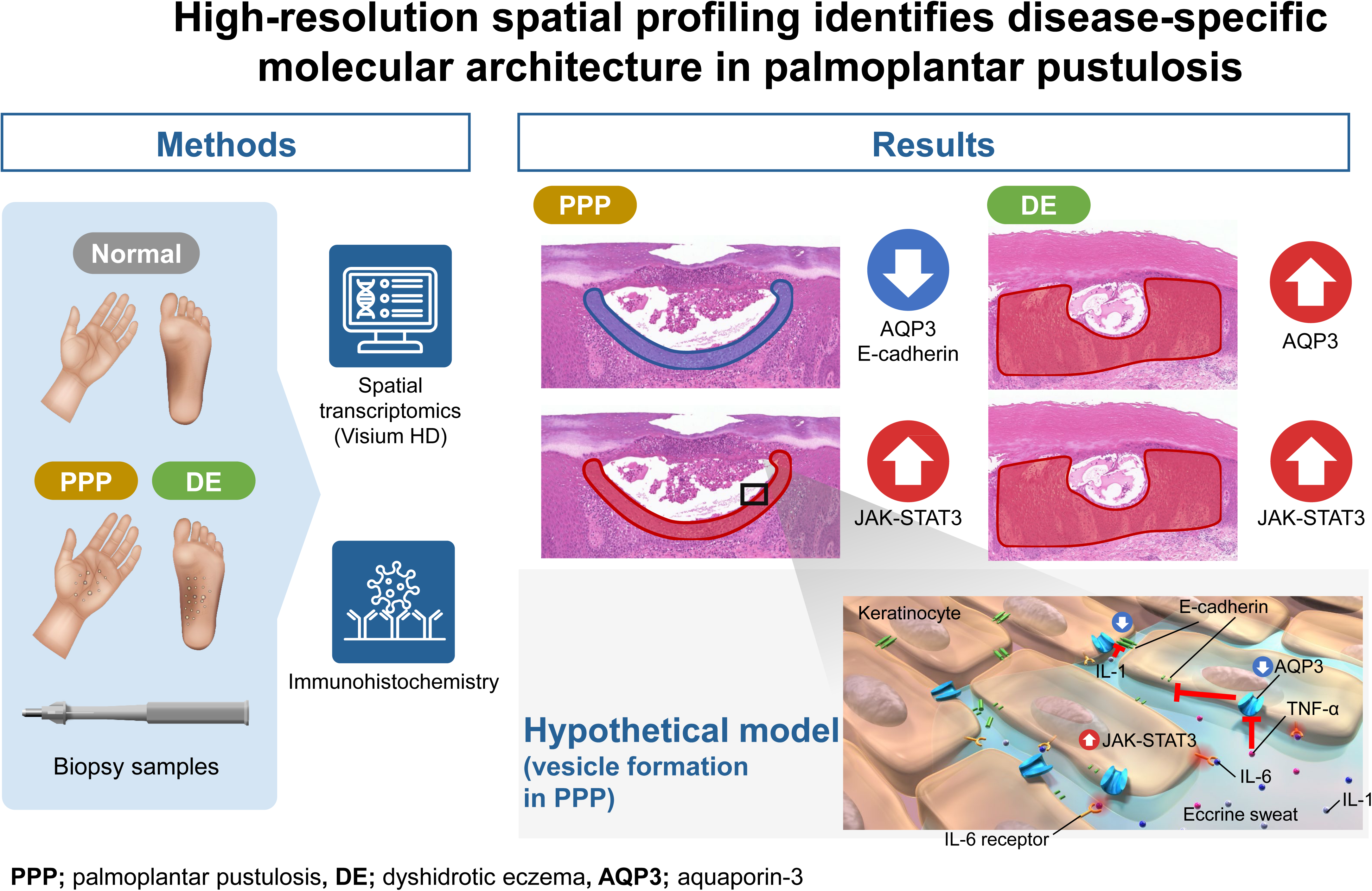

## Introduction

Palmoplantar pustulosis (PPP) and dyshidrotic eczema (DE) are chronic, treatment-refractory dermatoses characterized by intraepidermal vesiculation, resulting in marked impairment of the quality of life of patients. In this study, PPP is specifically defined as type-A PPP (the predominant form in Japan), which progresses from vesicles to pustules without psoriatic plaques (Hsu and Tsai, 2025; Murakami and Terui, 2020). In both conditions, lesions are generally confined to the palms and soles, making clinical distinction often challenging. Moreover, their underlying mechanisms remain poorly understood. Consequently, there are limited approved therapeutic options for these diseases, emphasizing a substantial unmet clinical need. Nevertheless, recent evidence has underscored fundamental differences in pathogenesis between PPP and DE.

In 2019, through our comparative analyses of hyaluronan levels in lesional epidermis, we reported that intraepidermal vesicles in PPP differed histologically from eczematous vesicles in DE (Murakami et al, 2019). In 2024, we further demonstrated that E-cadherin expression was reduced in keratinocytes surrounding PPP pustulo-vesicles and that IL-1 stimulation could reproduce this reduction in cultured keratinocytes (Yatsuzuka et al, 2024). In murine toe epidermis, sweat leakage and subsequent vesicle formation were detected using our custom fluorescent dye cocktail (JSAC) combined with two-photon excitation microscopy (Yatsuzuka et al, 2024). These results support our hypothesis that IL-1-rich eccrine sweat leaking from intraepidermal ducts triggers keratinocyte E-cadherin downregulation, inducing vesicle formation in PPP (Yatsuzuka et al, 2024). These findings strongly suggest that the mechanisms underlying vesicle formation in PPP are entirely distinct from those in DE. Around the same time, van Straalen et al. discovered through bulk RNA sequencing of lesional tissue that PPP exhibits an immune milieu resembling that of psoriasis, whereas DE, similar to atopic dermatitis, is dominated by Th2-driven inflammation (van Straalen et al, 2024). Altogether, these studies indicate that although PPP and DE appear clinically similar, their pathogenic foundations differ profoundly.

Considering that preventing intraepidermal vesicle formation may represent a curative approach, a detailed comparative investigation of the initiating mechanisms in PPP and DE is particularly important. Bulk RNA sequencing studies, such as that conducted by van Straalen et al. (van Straalen et al, 2024), are constrained by their inability to resolve microanatomical events within the highly organized three-dimensional architecture of the skin. More recently, Do et al. performed a spatial transcriptomic analysis of PPP; however, their study was limited to one patient (Do et al, 2025). Subsequently, Lee et al. expanded the spatial and single-cell characterization of PPP and identified a dermal immune niche marked by myeloid–lymphoid crosstalk and neutrophil-recruiting chemokine programs (Lee et al, 2026). Nonetheless, their study primarily addressed the immune–stromal architecture underlying pustular inflammation, rather than the epidermal programs that directly initiate the formation of intraepidermal vesicles. Therefore, in the present study, through spatial transcriptomic profiling of lesional tissues, we aimed to clarify the cellular programs underlying vesicle formation in PPP (*n* = 3) and DE (*n* = 3) and establish a framework for developing disease-specific curative therapies.

## Results

### Identification of lesion-associated epidermal clusters in PPP and DE

Figure 1a illustrates an overview of the spatial transcriptomic workflow. As described in the Methods section, the analysis included well-defined lesional samples of PPP (*n* = 3) and DE (*n* = 3), along with a normal palmar skin sample (*n* = 1) (Figure 1b–d). Through the integrated clustering of all specimens, 21 distinct epidermal and dermal clusters were identified (Figure 2a and b). Considering that the principal aim of this study was to clarify the mechanisms underlying intraepidermal vesicle formation, our subsequent analyses focused on epidermal clusters. In normal skin, layer-specific annotation was validated using canonical differentiation markers such as *LOR*, *KRT1*, *KRT2*, *KRT5*, *KRT10*, and *KRT14* (Figure 2c and d). Remarkably, both PPP and DE lesions contained a keratinocyte cluster (cluster 10) that was localized adjacent to vesicles (Figure 2b). Hence, this cluster was selected for downstream differential expression and pathway analyses.

**Figure 1.**
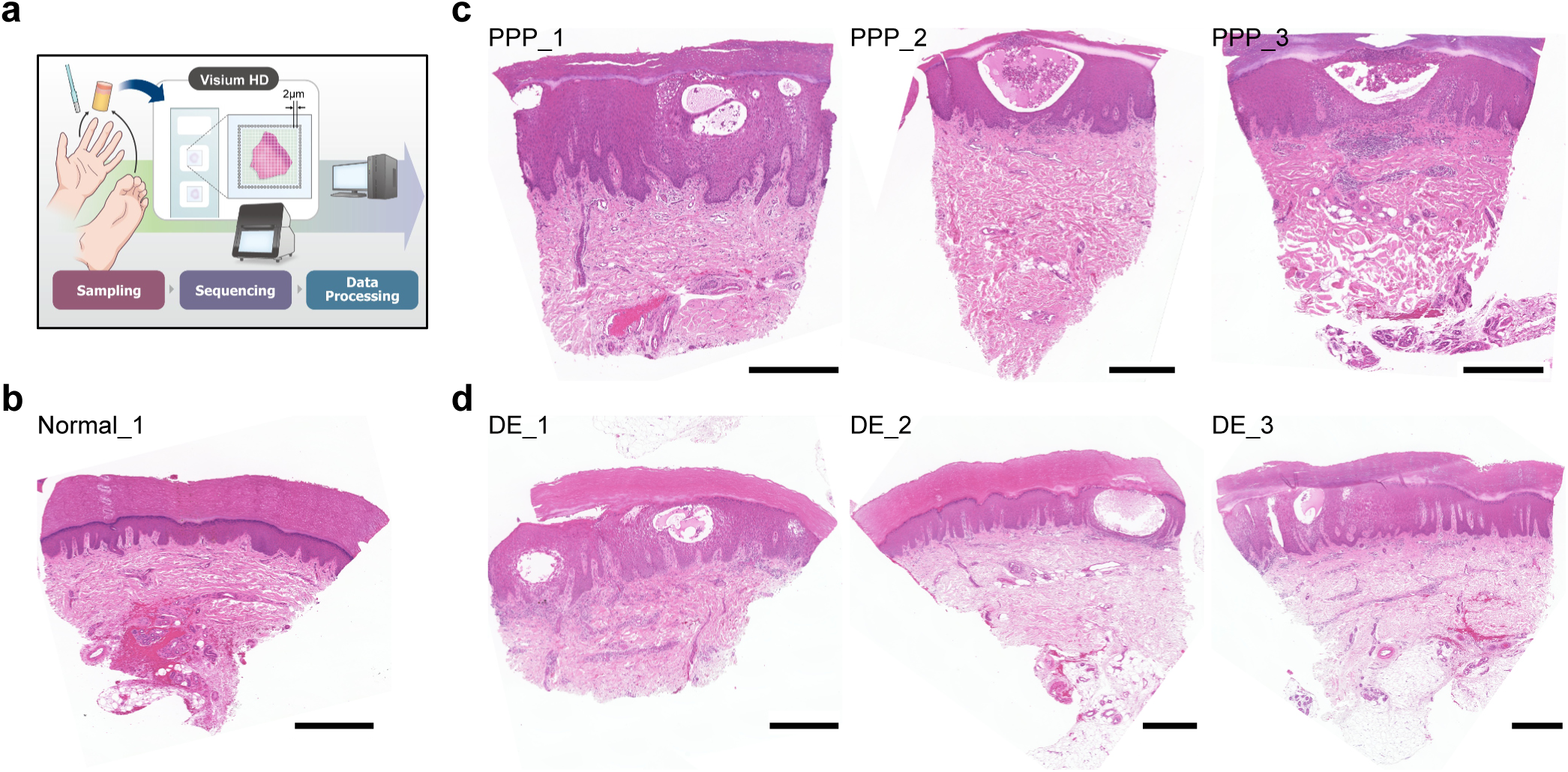
Experimental workflow and tissue specimens examined in this study. **(a)** Diagram showing the experimental workflow (© MEDICAL FIG). **(b–d)** H&E-stained sections of skin tissues: normal (b), palmoplantar pustulosis (PPP) (c), and dyshidrotic eczema (DE) (d). Scale bars, 500 μm.

**Figure 2.**
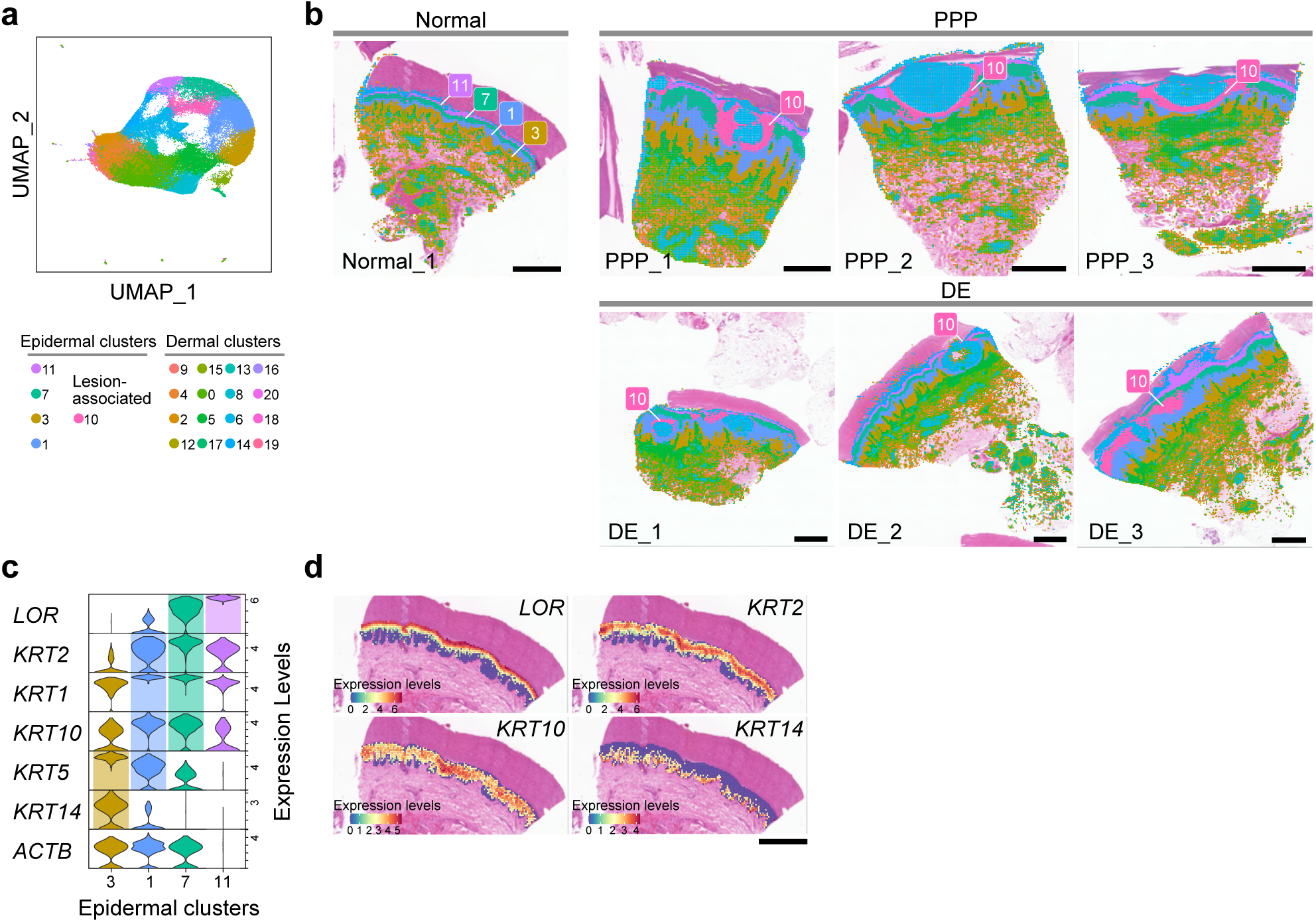
Stratification of skin tissue clusters and identification of lesion-associated subpopulations in PPP and DE specimens. **(a)** Uniform manifold approximation and projection (UMAP) plot of spatial clusters identified across normal, PPP, and DE samples. Epidermal clusters are labeled on the plot. **(b)** Spatial distribution of clusters overlaid on H&E-stained tissue images, with epidermal clusters labeled. **(c)** Violin plots depicting the expression of epidermal layer–specific marker genes in the indicated clusters. Cluster-matched colors indicate increased expression. *ACTB* is shown as a reference. **(d)** Spatial feature plots depicting the distribution of epidermal layer–specific genes in normal skin. Color scales represent gene expression levels.

### Validation of previously reported bulk RNA sequencing through spatial transcriptomic analysis

Our differential expression analysis revealed gene signatures consistent with the bulk RNA sequencing data reported by van Straalen et al. (van Straalen et al, 2024), wherein PPP and DE were defined using inclusion criteria widely comparable to those used in our study (as explained in the Methods section). In PPP lesions, *ABCA12*, *CXCL8*, *IL36A*, and *IL36G* were markedly upregulated (Figure 3a–c). Overall, these findings support the reproducibility of our analysis and confirm and extend previous bulk RNA sequencing results (van Straalen et al, 2024), incorporating spatial information that further highlights the vital role of the IL-36 pathway in PPP pathogenesis.

**Figure 3.**
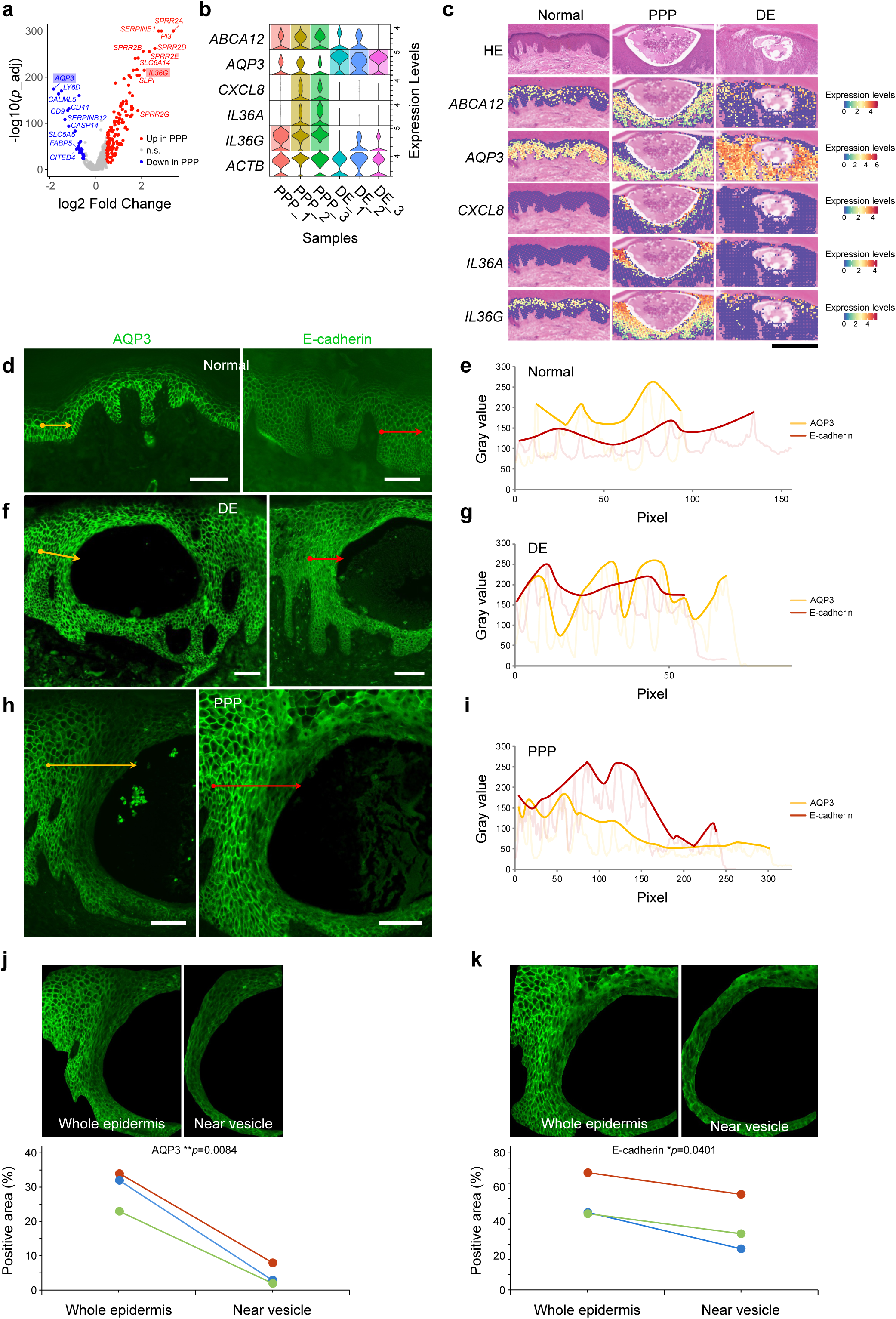
Decreased expression of aquaporin-3 (AQP3) and E-cadherin in keratinocytes adjacent to vesicles in PPP specimens. **(a)** Volcano plot depicting differentially expressed genes in lesion-associated clusters from PPP specimens compared with those from DE specimens. Genes upregulated (log_2_ fold change >0.5) and downregulated (log_2_ fold change <−0.5) in PPP are depicted in red and blue, respectively. Genes with more than twofold changes (log_2_ fold change >1 or <−1) are labeled, with *AQP3* and *IL36G* highlighted in blue and red, respectively. Adjusted *p* values (p_adj) were calculated using the Benjamini–Hochberg method. **(b)** Violin plots illustrating the expression of upregulated and downregulated genes associated with PPP in lesion-related clusters. Increased expression levels are highlighted with sample-matched colors. *ACTB* is included as a reference. **(c)** Spatial feature plots displaying differentially expressed genes distribution between PPP and DE specimens. Corresponding H&E-stained images of the analyzed areas are shown above. Color scales represent expression levels. **(d–i)** Representative immunofluorescence images and corresponding fluorescence intensity line-profile graphs of normal skin (d, e), DE lesions (f, g), and PPP lesions (h, i). In each image panel, a line drawn from a circular starting point to an arrowhead endpoint indicates the region used for fluorescence intensity profile analysis. In each graph (e, g, and i), the horizontal axis indicates pixel distance along the drawn line, and the vertical axis indicates fluorescence intensity. Pale lines represent the raw fluorescence intensity profiles, and bold lines represent membrane-associated signal profiles generated by extracting and connecting peak intensities. **(j and k)** Quantification of immunofluorescence staining in PPP lesions. The upper panels depict representative images illustrating the region of interest (ROI) used for analysis. The lower panels depict the quantified positive area for AQP3 (j) and E-cadherin (k). Each line represents one PPP sample (n = 3). Statistical significance was evaluated using a two-tailed paired Student’s *t*-test; \*\**p* < 0.01 and \**p* < 0.05. Scale bars, 100 μm.

### Reduced aquaporin-3 (AQP3) expression as a distinct feature of PPP

As the most striking observation of our study, the expression of *AQP3* in PPP cluster 10 was considerably reduced compared with that in DE and normal epidermis (Figure 3a–c). Conversely, DE showed markedly increased *AQP3* expression throughout the epidermis, including in keratinocytes adjacent to vesicles (Figure 3c). To validate these findings at the protein level, we conducted immunofluorescence staining for AQP3 in PPP, DE, and normal skin samples (Figure 3d–k) and, in parallel, examined the expression of E-cadherin, as previous studies have demonstrated a close association between AQP3 and E-cadherin in intercellular adhesion among epidermal keratinocytes (Blaydon and Kelsell, 2014; Kim and Lee, 2010). Furthermore, to visualize the spatial pattern of AQP3 and E-cadherin expression, we generated fluorescence intensity line profiles from representative images (Figure 3d–i). These line-profile analyses demonstrated that in normal and DE epidermis, the fluorescence intensities of both AQP3 and E-cadherin remained relatively uniform along the epidermal layer (Figure 3d–g). Conversely, in PPP epidermis, reduced fluorescence intensity of both proteins was detected in regions adjacent to intraepidermal vesicles (Figure 3h and i), suggesting a localized loss of AQP3 and E-cadherin in keratinocytes surrounding vesicles.

To further validate these findings in PPP, we additionally conducted a quantitative region of interest (ROI)-based analysis of PPP lesions. For AQP3, the fluorescence-positive area was consistently lower in the near-vesicle region than in the whole-epidermal region, and this reduction reached statistical significance (*p* = 0.0084, Figure 3j). Similarly, E-cadherin expression was also reduced in keratinocytes in near-vesicle regions compared with that in whole-epidermal regions, and this difference was statistically significant (*p* = 0.0401, Figure 3k). These protein-level findings were consistent with the spatial expression pattern observed in the transcriptomic data. Altogether, these findings suggest that localized loss of AQP3 and E-cadherin is associated with keratinocyte dysfunction in vesicle-adjacent epidermis in PPP. Due to the limited availability of DE specimens suitable for ROI-based quantification of AQP3/E-cadherin immunofluorescence, ROI-based quantitative analysis was not performed for DE in the present study. Nevertheless, unlike PPP, DE has been reported to exhibit increased AQP3 expression (Soler et al, 2015).

### Spatially differential activation of JAK-STAT3 signaling in the epidermis between PPP and DE

Pathway enrichment analysis revealed further disease-specific signatures. In PPP cluster 10, JAK-STAT3 signaling was strongly activated (Figure 4a). Moreover, single-sample gene set enrichment analysis (ssGSEA) indicated that JAK-STAT3 signaling was activated in DE but was distributed across the entire epidermis rather than in vesicle-adjacent keratinocytes alone (Figure 4b). Examination of the downstream targets of STAT3 confirmed the expression levels of *CXCL1* and *SOCS3* in PPP cluster 10 (Figure 4c and d). These findings were corroborated by immunohistochemistry (Figure 4e–i), which revealed intense phosphorylated STAT3 (p-STAT3) nuclear staining in keratinocytes immediately beside the vesicles in PPP (Figure 4e and h). In DE, the expression of p-STAT3 was more uniform (Figure 4f and i). These findings are consistent with those reported by van Straalen et al. on STAT3 as a central node in PPP pathogenesis (van Straalen et al, 2024) and also emphasize the importance of spatially concentrated activation in vesicle formation. To further strengthen these findings, we performed additional quantitative image analysis to examine p-STAT3 immunoreactivity in regions adjacent to vesicles and in more distant epidermal areas (Figure 4j–m). In PPP lesions, p-STAT3 3,3′-diaminobenzidine (DAB) intensity was significantly higher in epidermal keratinocytes adjacent to the vesicle than in keratinocytes located farther away (*p* = 0.0408, Figure 4j and k). This pattern was consistently detected across the three PPP cases (Figure 4k), indicating the localized enhancement of STAT3 activation in near-vesicle epidermis. In contrast, we found no significant difference in p-STAT3 DAB intensity between near and far epidermal keratinocytes in DE lesions (*p* = 0.4770, Figure 4l and m). Although case-to-case variation was present, the paired analysis did not support a reproducible near-vesicle increase in p-STAT3 expression in DE. Overall, these results suggest that localized activation of STAT3 signaling in keratinocytes adjacent to vesicular structures is characteristic of PPP but not observed in DE.

**Figure 4.**
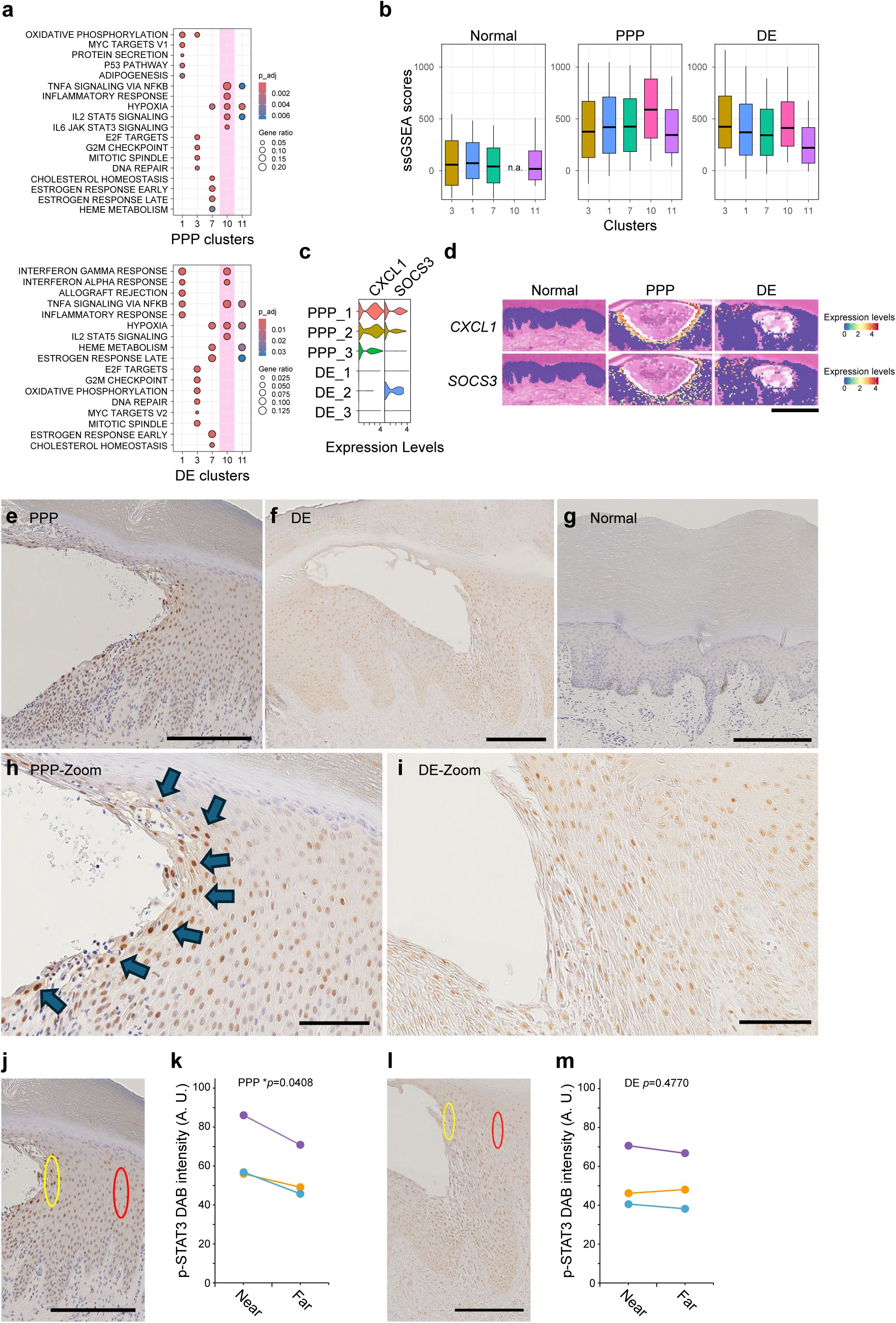
Distinct activation patterns of the JAK-STAT3-mediated pathway in lesion-associated areas of PPP and DE specimens. **(a)** Dot plots representing the enriched MSigDB hallmark gene sets in the epidermal clusters of PPP (upper panel) and DE (lower panel) samples. The top five hallmark gene sets enriched in each cluster are shown. Gene ratio represents the proportion of input genes linked to each term. The color scale denotes the adjusted *p* value (p_adj), representing statistical significance after multiple testing correction (Benjamini–Hochberg method). **(b)** Box plots showing single-sample gene set enrichment analysis (ssGSEA) scores for the IL6-JAK-STAT3 signaling pathway in epidermal clusters. **(c)** Violin plots showing the expression levels of known STAT3 target genes in PPP and DE samples. **(d)** Spatial feature plots depicting the expression patterns of the STAT3 target genes within the epidermis. **(e–i)** Representative chromogenic immunohistochemistry images of phosphorylated STAT3 (p-STAT3) in PPP (e, h: representative of 3), DE (f, i: representative of 3), and normal skin (g). Arrows in (h) indicate keratinocyte nuclei showing particularly intense p-STAT3 staining. **(j–m)** Quantitative analysis of p-STAT3 immunohistochemistry. (j) and (l) depict representative images illustrating the regions selected for quantification in PPP and DE specimens, respectively. Yellow and red ellipses indicate the near-vesicle and far-vesicle ROIs, respectively. (k) and (m) show the quantified p-STAT3 3,3′-diaminobenzidine (DAB) signal intensity in PPP and DE specimens, respectively. Mean DAB intensity was measured in near-vesicle and far-vesicle epidermal regions and is presented in arbitrary units (A.U.). Each line represents one sample (n = 3 per group). Statistical significance was evaluated using a two-tailed paired Student’s *t*-test; \**p* < 0.05. Scale bars, 250 µm (d, e, f, g, j, l) and 100 µm (h, i).

## Discussion

This study provides high-resolution spatial transcriptomic evidence that delineates epidermal transcriptional signatures unique to PPP and DE. Consistent with our aim to clarify the cellular programs underlying vesicle formation in PPP and DE, the identification of a keratinocyte cluster (cluster 10) localized adjacent to vesicles is particularly significant. Although both diseases exhibit inflammatory activation of JAK-STAT3 signaling, they differ markedly in microanatomical localization and upstream molecular context.

Xiaoling et al. demonstrated robust IL-36G expression in keratinocytes adjacent to pustules in PPP, suggesting that this protein is involved in neutrophil recruitment and pustule development (Xiaoling et al, 2019). Our spatial data reinforce this concept and, combined with the findings of van Straalen et al. (van Straalen et al, 2024), emphasize IL-36A as an additional contributor (Figure 3b and c). A phase IIb trial of spesolimab, an anti-IL-36 receptor antibody, enrolled a substantial number of Japanese patients with PPP and investigated the responses in the wider Asian subgroup; however, its primary endpoint was not met (Burden et al, 2023). The considerable placebo response and the inherently fluctuating, self-remitting nature of PPP may have contributed to the lack of efficacy, thereby obscuring actual treatment effects. Furthermore, the relatively small sample size and the short observation period (16 weeks) may have restricted the statistical power required to establish a clear therapeutic benefit of spesolimab in comparison with the placebo (Burden et al, 2023). Hence, this outcome does not necessarily indicate that IL-36 is not involved in the pathogenesis of PPP; further investigation is necessary. In PPP lesions, the expression levels of AQP3 and E-cadherin were reduced in a keratinocyte cluster (cluster 10) localized adjacent to vesicles, whereas in DE lesions, the expression was preserved or diffuse (Figure 3), indicating a potentially unique mechanism underlying vesicle formation in PPP. AQP3, a transporter of water and glycerol expressed throughout the epidermis, helps in maintaining E-cadherin-mediated adhesion (Blaydon and Kelsell, 2014). It has been reported that the surface expression of E-cadherin decreases through siRNA-mediated AQP3 knockdown, indicating that AQP3 is not only a passive water channel but also a key regulator of keratinocyte cohesion (Kim and Lee, 2010). These findings suggest that reduced AQP3 expression may represent an upstream event that predisposes keratinocytes to IL-1-mediated E-cadherin downregulation and vesicle formation in PPP. Considering that human eccrine sweat contains multiple cytokines, including TNF-α (Baker, 2019), which itself reduces AQP3 expression in keratinocytes via p38/ERK activation (Horie et al, 2009), a dual hit, namely, TNF-α-driven AQP3 loss and IL-1-induced E-cadherin suppression, probably underlies vesicle formation in PPP. Therefore, therapeutic strategies that preserve or restore keratinocyte AQP3 may be uniquely effective in PPP. In fact, agents such as retinoic acid (Bellemere et al, 2008) and SIG-1191 (N-acetylglutaminoyl-S-farnesyl-L-cysteine) (Fernandez et al, 2017) have been demonstrated to improve AQP3 expression. In particular, SIG-1191 has already been investigated as a cosmetic ingredient. It is necessary to explore the effects of these substances on PPP-related skin symptoms when investigating treatment options.

Our results also suggest that JAK-STAT3 signaling is a shared therapeutic target in PPP and DE (Figure 4). In clinical trials, delgocitinib cream, a topical pan-JAK inhibitor, has already been demonstrated to be effective in treating chronic hand eczema, including DE (Bissonnette et al, 2024). Hence, systemic JAK inhibitors may be highly effective as treatment options for refractory or severe cases. Regarding PPP, multiple case reports and case series from diverse regions and ethnic groups have demonstrated that systemic JAK inhibitors such as tofacitinib and upadacitinib are also effective (Gleeson et al, 2023; Huang et al, 2025; Taniguchi and Yamamoto, 2024; Yang et al, 2025), thereby strengthening further expectations. Prospective clinical trials of systemic JAK inhibition specifically in type-A PPP are warranted to substantiate these expectations. Nevertheless, although rare, PPP-like eruptions associated with JAK inhibitors have been reported (Shibata et al, 2019). In Japan, apremilast was approved for PPP treatment in 2025 (Terui et al, 2025a). Apremilast may modulate JAK-STAT3 signaling through the cAMP-PKA-CREB-SOCS3 pathway (Li et al, 2019). Our findings further support this potential mechanism. In particular, our study showed that STAT3 activation in PPP is particularly prominent in a keratinocyte cluster (cluster 10) adjacent to vesicles, suggesting that modulation of the JAK-STAT3 pathway in this disease can potentially target intraepidermal vesicle formation itself directly. Considering that human eccrine sweat also contains IL-6 (Baker, 2019) and that IL-6 can activate JAK-STAT3 signaling in epidermal keratinocytes (Liu et al, 2021; Piao et al, 2019), the strongest activation occurring in keratinocytes beside vesicles in PPP specimens, where eccrine sweat can possibly leak into the epidermis, is reasonable (Yatsuzuka et al, 2024). This observation supports our hypothesis (see Graphical abstract).

Despite the insights gained, this study has several limitations. A major limitation is the modest sample size. Increasing the sample size was constrained by the high cost of high-resolution spatial transcriptomics platforms, such as Visium HD. Remarkably, PPP has recently been recognized to demonstrate highly heterogeneous immune microenvironments on a case-by-case basis (Terui et al, 2025b). Moreover, its pathophysiology undergoes rapid changes depending on the disease stage, ranging from the vesicle phase to the pustulo-vesicle phase and then to the pustule phase. Accordingly, using a more diverse set of specimens and conducting large-scale analyses in future studies are necessary to capture this complexity. Another limitation is that we discussed the key hypothesis we established, that is, the downregulation of AQP3 expression acts upstream of reduced E-cadherin expression, on the basis of immunohistochemical findings and previous literature. Nevertheless, our study lacks original in vitro experimental data that could directly demonstrate this causal relationship. As a future direction, this hypothesis must be explored by investigating how the siRNA-mediated knockdown or pharmacological inhibition of AQP3 affects the surface expression of E-cadherin, in both cultured epidermal keratinocytes and three-dimensional models of cultured epidermis.

In conclusion, our spatial transcriptomic analysis delineated the epidermal signatures that are not only distinct but also partially overlap between PPP and DE. JAK-STAT3 signaling was activated in both PPP and DE, suggesting shared inflammatory pathways between these two diseases. Nonetheless, compared with DE, PPP exhibited focal reduction of AQP3 and E-cadherin expression in keratinocytes adjacent to vesicles, accompanied with markedly increased activation of JAK-STAT3 signaling in these cells. These results may provide important insights into the mechanisms underlying vesicle formation in PPP. Therapeutic strategies targeting JAK-STAT3 pathways may benefit both PPP and DE, and those intended to restore AQP3-mediated adhesion could be particularly relevant to PPP.

## Materials & Methods

### Sex as a biological variable

Because PPP is more prevalent among females in the Japanese population, only female patients with PPP were included in this study. Therefore, the potential effects of sex on disease-associated molecular signatures could not be evaluated.

### Patients and skin samples

The histopathological definitions of PPP and DE in this study were based on the diagnostic histopathological features proposed by Masuda-Kuroki et al. in 2019 (Masuda-Kuroki et al, 2019). Briefly, PPP referred to vesicles without spongiosis, accompanied with microabscesses at the edges of these vesicles. In contrast, vesicles with spongiosis but without microabscesses at their edges defined DE. As this study included only type-A PPP cases, patients with psoriatic plaques were excluded from the analysis. In the spatial transcriptomic analysis, we used 2–4 mm punch biopsy specimens obtained from six patients with PPP or DE (three patients in each disease group, Table 1) treated at Ehime University Hospital. Immunohistochemical analysis was conducted using an independent cohort, different from the patients included in the spatial transcriptomic analysis. Immunofluorescence staining for AQP3 and E-cadherin and chromogenic immunohistochemistry for p-STAT3 were performed using separate validation sample sets. In all analyses, normal skin on the palm or sole adjacent to the nevus was used as the control.

**Table 1.** Clinical data from biopsied patients used for spatial profiling.

| **PPP** | **Patient 1** | **Patient 2** | **Patient 3** |
| --- | --- | --- | --- |
| Age (year) | 53 | 50 | 39 |
| Sex | Female | Female | Female |
| Anatomical site, biopsied | Palm (2 mm) | Sole (2 mm) | Sole (2 mm) |
| Phase, biopsied | Pustulo-vesicle | Pustulo-vesicle | Pustulo-vesicle |
| Systemic treatment | None | None | None |
| Smoking status | Never | Past | Current |
| Shown as | PPP_1 | PPP_2 | PPP_3 |
| **DE** | **Patient 1** | **Patient 2** | **Patient 3** |
| Age (year) | 56 | 76 | 33 |
| Sex | Male | Male | Female |
| Anatomical site, biopsied | Sole (3 mm) | Palm (3 mm) | Palm (4 mm) |
| Phase, biopsied | Vesicle | Vesicle | Vesicle |
| Systemic treatment | Antihistamine | Antihistamine | None |
| Smoking status | Never | Never | NA |
| Shown as | DE_1 | DE_2 | DE_3 |

PPP: palmoplantar pustulosis, DE: dyshidrotic eczema, NA: not available. Not available indicates data were not recorded in the medical record.

### Spatial transcriptomics

Spatial gene expression in FFPE tissue sections was profiled using the Visium HD Spatial Gene Expression Reagent Kit (10× Genomics, PN-1000668), according to the manufacturer’s instructions. Briefly, 5-μm-thick FFPE tissue sections were incubated overnight at 50°C with predefined pairs of gene-specific probes (Visium Human Transcriptome Probes v2, 10× Genomics, PN-1000466), followed by probe ligation at 37°C for 60 min. Single-stranded ligation products were released from the tissue and captured by spatially barcoded capture probes within the Visium HD slide Capture Area. We extended the captured probes to incorporate the spatial barcode and unique molecular identifier (UMI). Libraries were prepared using the Visium HD library reagents (10× Genomics, PN-1000668) with a Visium CytAssist (10× Genomics), followed by sequencing on the Illumina NovaSeq 6000 platform. Using Space Ranger v3.0 (10× Genomics), we aligned raw sequencing reads to the human reference genome (GRCh38, version 2020-A, 10× Genomics), generating UMI count matrices with spot-associated barcodes.

### Analysis of spatial transcriptomics data

UMI count matrices at a 16-µm bin resolution from all samples (normal, PPP_1, PPP_2, PPP_3, DE_1, DE_2, and DE_3) were imported into Seurat (v5.3.0) (Hao et al, 2024) running on R (v4.4.2). After merging, we excluded spots with <10 detected genes. After log-normalization of gene expression values using the NormalizeData() function, we reduced the dimensionality using RunPCA() and integrated the datasets using the Harmony algorithm. Specimens were then clustered using the FindNeighbors() and FindClusters() functions (dims = 1:11, resolution = 0.55), and results were visualized in uniform manifold approximation and projection (UMAP) space using RunUMAP(). Subsequently, epidermal clusters, including lesion-associated subsets, were extracted for downstream analysis.

After identifying differentially expressed genes between PPP and DE samples using FindAllMarkers(), we calculated the *p* values using the Wilcoxon test and adjusted them using the Benjamini–Hochberg method. Moreover, gene expression levels were visualized using violin plots and mapped onto tissue sections using SpatialFeaturePlot(). ggplot2 (v3.5.2) was used for generating box plots and volcano plots. Next, clusterProfiler (v4.14.6) was used to perform threshold-based pathway enrichment analysis using cluster marker genes identified using Seurat and Hallmark gene sets obtained from msigdbr (v25.1.1). In addition, using the escape package (v2.2.3), we conducted single-sample GSEA to calculate MSigDB hallmark pathway activity scores within epidermal clusters from PPP and DE samples.

### Immunofluorescence and chromogenic immunohistochemistry of human skin samples

FFPE specimens were prepared for the immunohistochemical analysis of each target molecule in the lesional epidermis. For immunofluorescence staining, 5-μm-thick tissue sections were deparaffinized and rehydrated for heat-mediated antigen retrieval. The sections were then incubated overnight at 4°C in a humidified chamber with primary antibodies (anti-AQP3 [1:500; Abcam, ab125219] or anti-E-cadherin [1:500; Abcam, ab40772]) diluted in a background-reducing antibody diluent (Agilent, S3022). An Alexa Fluor 488–conjugated goat anti-rabbit IgG (1:200) was used as the secondary antibody. For chromogenic immunohistochemistry, we incubated the sections with a primary antibody (anti-p-STAT3 [1:200; Cell Signaling Technology, 9145]) and then treated them with a horseradish peroxidase–conjugated secondary antibody. These sections were visualized using DAB as the chromogen.

### Quantification of immunofluorescence staining

Immunofluorescence images were acquired using a BZ-X710 fluorescence microscope (Keyence). Image analysis was conducted using the BZ-X Analyzer software (Keyence). As an initial quantitative assessment of signal distribution, fluorescence intensity line profiles were generated for representative images from normal epidermis, DE lesion, and PPP lesion. Using the line-profile function in BZ-X Analyzer, a linear ROI, defined from a circular starting point to an arrowhead endpoint, was manually drawn across the epidermal layer, and the entire length of the ROI was examined, including areas adjacent to intraepidermal vesicles in DE and PPP samples. Fluorescence intensity values (gray values) along the line were plotted against pixel distance to visualize spatial changes in signal intensity for AQP3 and E-cadherin. Because both AQP3 and E-cadherin are membrane proteins, the original line-scan data were treated as raw profiles, and, to more clearly illustrate the visual trend, an additional membrane-associated signal profile was generated by extracting and connecting only the peak intensities. For further quantitative analysis, the fluorescence images of PPP lesions (*n* = 3) were thresholded manually to detect AQP3- or E-cadherin-positive signals. The same threshold settings were applied to all images within each staining condition to ensure consistent quantification. ROIs were manually defined for two epidermal compartments, viz., the whole epidermis and the epidermal region adjacent to intraepidermal vesicles, defined as the five keratinocyte layers from the vesicle side (near vesicle). Within each ROI, the percentage of positive area for AQP3 and E-cadherin was quantified using the Hybrid Cell Count and Macro Cell Count functions of the BZ-X Analyzer software. Three independent biopsy samples obtained from patients with PPP were examined. For each sample, measurements obtained from the whole epidermis and the near-vesicle epidermis were treated as paired observations.

### Quantification of DAB signal

Quantification of p-STAT3 immunohistochemical staining was performed using ImageJ (Fiji distribution). ROIs were manually defined within the epidermis using standardized elliptical selections with a fixed size (width: 40 μm; height: 140 μm) to ensure consistency across samples. Two ROIs were defined for each specimen, viz., a near-vesicle ROI (near), positioned adjacent to the vesicle cavity and encompassing approximately the first three layers of epidermal keratinocytes immediately surrounding the vesicle wall, and a far-vesicle ROI (far), placed within the same epidermal layer but at a sufficient distance from the near-vesicle ROI in the outward direction. For each ROI, the mean pixel intensity of the DAB signal corresponding to p-STAT3 immunoreactivity was quantified using the standard measurement functions in ImageJ. Signal intensity was expressed in arbitrary units (A.U.). Three independent biopsy samples obtained from patients with PPP and DE (n = 3 each) were analyzed.

### Statistics

Data are expressed as mean ± SD. Statistical analyses were conducted according to the type of data and comparison being evaluated. For comparisons between two groups in the spatial transcriptomic analysis, the Wilcoxon test was used to calculate *p* values and determine statistical significance. Next, we adjusted the resulting *p* values for multiple testing using the Benjamini–Hochberg method. For the quantification of immunofluorescence staining or DAB signal, statistical analyses were conducted using paired comparisons within the same specimen. To compare the fluorescence-positive area between the whole epidermis and the near-vesicle region, as well as the mean pixel intensity of the DAB signal between the near-vesicle and far-vesicle regions, a two-tailed paired Student’s *t*-test was used. *p* < 0.05 was considered statistically significant.

### Ethics Statement

This study was performed in accordance with the Declaration of Helsinki. Collection of human tissue samples for this study was approved as part of the study protocol. This human study was approved by the Institutional Ethics Committee of Ehime University School of Medicine - approval: 1802009, 1808003, and 2501007. All adult participants provided written informed consent to participate in this study.

## Abbreviations

PPP: palmoplantar pustulosis
DE: dyshidrotic eczema
AQP3: aquaporin-3
ROI: region of interest
ssGSEA: single-sample gene set enrichment analysis
p-STAT3: phosphorylated STAT3
DAB: 3,3′-diaminobenzidine
UMI: unique molecular identifier
UMAP: uniform manifold approximation and projection.

## Data Availability Statement

The spatial transcriptomics data discussed in this publication are available in the Gene Expression Omnibus database (accession no. GSE310984) of NCBI. Additional processed data supporting the findings of this study are available from the corresponding author upon reasonable request.

## Acknowledgments

This study was partially supported by JSPS KAKENHI (grant number 23K07768), the UCB Research Grant Award 2025 from the Japanese Society for Psoriasis Research, and a research grant from Sun Pharma Co. Ltd. We thank Prof. Tasuku Nishihara, Department of Anesthesia and Perioperative Medicine, Ehime University Graduate School of Medicine, Ehime, Japan, for his technical support. We also thank Takahiro Fukazawa and Taro Kuroda of the Division of Medical Research Support, Advanced Research Support Center (ADRES), Ehime University, Ehime, Japan, for their technical support. We are grateful to Enago (www.enago.jp) for the English language review and to the study participants for their time and for providing biopsy samples.

## Author Contributions Statement

Conceptualization: KY, JM, MaM; Data Curation: KY, YM, KI, DS, MeM; Formal Analysis: KY, YM, KI, DS, MeM, YF; Funding Acquisition: KY, MaM; Investigation: KY, YM, KI, DS, MeM, TT, KS; Project Administration: KY; Resources: KY, KS, MaM; Supervision: YF, MaM; Visualization: KY, YM, KI, DS, MeM, MaM; Writing — Original Draft Preparation: KY, MaM; Writing — Review and Editing: KY, JM, YM, KI, DS, MeM, TT, KS, YF, MaM.

## Declaration of Generative AI use

During the preparation of this work, the authors used ChatGPT to improve the fluency and clarity of their English writing. After using this tool, the authors reviewed and edited the content as needed and took complete responsibility for the publication’s content.

